# Analysis of Longitudinal Change Patterns in Developing Brain Using Functional and Structural Magnetic Resonance Imaging via Multimodal Fusion

**DOI:** 10.1101/2024.04.07.588473

**Authors:** Rekha Saha, Debbrata K. Saha, Zening Fu, Marlena Duda, Rogers F. Silva, Vince D. Calhoun

## Abstract

Functional and structural magnetic resonance imaging (fMRI and sMRI) are complementary approaches that can be used to study longitudinal brain changes in adolescents. Each individual modality offers distinct insights into the brain. Each individual modality may overlook crucial aspects of brain analysis. By combining them, we can uncover hidden brain connections and gain a more comprehensive understanding. In previous work, we identified multivariate patterns of change in whole-brain function during adolescence. In this work, we focus on linking functional change patterns (FCPs) to brain structure. We introduce two approaches and applied them to data from the Adolescent Brain and Cognitive Development (ABCD) dataset. First, we evaluate voxelwise sMRI-FCP coupling to identify structural patterns linked to our previously identified FCPs. Our approach revealed multiple interesting patterns in functional network connectivity (FNC) and gray matter volume (GMV) data that were linked to subject level variation. FCP components 2 and 4 exhibit extensive associations between their loadings and voxel-wise GMV data. Secondly, we leveraged a symmetric multimodal fusion technique called multiset canonical correlation analysis (mCCA) + joint independent component analysis (jICA). Using this approach, we identify structured FCPs such as one showing increased connectivity between visual and sensorimotor domains and decreased connectivity between sensorimotor and cognitive control domains, linked to structural change patterns (SCPs) including alterations in the bilateral sensorimotor cortex. Interestingly, females exhibit stronger coupling between brain functional and structural changes than males, highlighting sex-related differences. The combined results from both asymmetric and symmetric multimodal fusion methods underscore the intricate sex-specific nuances in neural dynamics. By utilizing two complementary multimodal approaches, our study enhances our understanding of the dynamic nature of brain connectivity and structure during the adolescent period, shedding light on the nuanced processes underlying adolescent brain development.

## 1. Introduction

The intricate neural network within the brain is recognized as one of the most complex systems in existence. A modern neuroscience approach involves viewing brain regional interactions as a graph network, known as the brain connectome or brain network [1]. By understanding network properties, researchers can gain insight into how the brain manages information flow by transferring neural signals among brain regions [2]. Magnetic resonance imaging (MRI) is a widely utilized method for obtaining comprehensive brain information and one of the only modalities that is capable of visualizing both structure and function. Structural neuroimaging modalities, such as sMRI and diffusion MRI, provide insights into the anatomical structure and tissue composition of the brain while functional neuroimaging modalities, such as fMRI based on bloodoxygenation-level-dependent signal, provide indirect measurements of brain function and activity [3, 4]. fMRI is a key method for assessing functional connectivity (FC) by analyzing temporal coherence among different brain regions using resting-state data. Most studies have still examine functional and structural measures independently when analyzing the brain. However there is considerable evidence that combining structural and functional MRI data can lead to benefits [5, 6]. Multimodal fusion in neuroimaging integrates information obtained from various imaging techniques, aiming to overcome individual modality limitations and gain deeper insights into brain dynamics [7, 8, 9]. The primary goal of multimodal fusion is to enhance the analytical capabilities of each modality through combined analysis, rather than treating each modality separately.

Prior research in multimodal analysis often involves examining data from distinct modalities separately and later combining the independent outcomes obtained from these individual analyses. Alternatively, some studies have used information from one modality to constrain or guide models related to another modality. While these approaches have been beneficial, they can under utilize cross-modality information. There is a growing trend in approaches that employ symmetric data fusion methods [7] to capitalize on joint information across modalities. Feature-based symmetric data fusion methods initially extract valuable, high-dimensional features from diverse modalities and then explores the connections among these features. This method effectively uses the complementary information present in each modality to reveal variations in data that might not be evident through unimodal analyses. Various studies have highlighted the potential of employing such cross-modality or joint information in understanding the human brain and its disorders. These methods have been instrumental in characterizing diseases, identifying potential biomarkers, and unraveling disrupted connections in complex mental illnesses [7].

Each multivariate fusion method possesses distinct optimization priorities and limitations. For instance, methods like mCCA [10] and partial least squares (PLS) [11, 12] facilitate both common and distinct levels of connection across modalities. However, these methods may not achieve adequate spatial sparsity in their separated sources. For instance, mCCA emphasizes inter-subject covariation across two feature sets, generating linked variables known as canonical variants (CVs). These CVs correlate with each other solely on the same indices, and their corresponding correlation values are termed canonical correlation coefficients (CCC). While this approach captures both common and distinct aspects of features, the resulting brain maps for multiple components may appear similar if the CCCs lack sufficient distinctiveness. On the other hand, spatial decomposition approaches like jICA [13] and linked independent component analysis [14] maximize independence among estimated sources combining multiple modalities, but they primarily allow a common mixing matrix. While these methods detect common features across all modalities effectively, they may neglect features distinct to one or more modalities, especially when combining more than two modalities. Several prior studies that merged function and structure [15, 16, 5, 17] support the idea that components derived from each modality exhibit some correlation in their mixing profiles among subjects. This serves as motivation to utilize an approach that aims for optimal inter-modal association flexibility while ensuring robust source separation capabilities.

In order to analyze the shared information among the features found in different imaging modalities, we used mCCA+jICA method, a widely recognized and extensively used multimodal fusion approach [18]. mCCA+jICA, is a data-driven multivariate fusion technique [19, 13] that simultaneously analyzes multimodal data by combining mCCA and jICA in a two-step process [7]. In the first step, mCCA is utilized to identify highly correlated components between multiple modalities [18, 20]. This is followed by the application of jICA in the second step to decompose these correlated components into spatially independent components, known as ICs. The mCCA+jICA algorithm has been employed [18] to combine fMRI contrast maps and diffusion tensor imaging (DTI) fractional anisotropy maps for examining group differences among healthy controls (HCs), schizophrenia patients, and bipolar patients. Importantly, that study found that the combined algorithm yielded increased accuracy in group classification compared to using the constituent algorithms individually. Ouyang et al. used mCCA+jICA to identify patterns of gray matter (GM) and white matter (WM) covariance in patients with Alzheimer’s disease [21]. Similarly, Kim et al. applied mCCA+jICA to analyze multimodal sMRI and DTI data from patients with obsessivecompulsive disorder and HCs, uncovering significant alterations in interconnected networks of GM and WM [22]. But, to the best of our knowledge, no prior studies have investigated the estimation of changes in multivariate pattern coupling in FNC and GMV associated with age progression, utilizing the mCCA+jICA multimodal fusion analysis method.

In our study, we present two innovative methodologies: 1) investigate the relationship between multivariate functional change patterns (FCPs) and voxel-wise gray matter (ΔGMV) data to explore within-subject age-related changes in whole-brain structure and function at an individual level 2) employed a symmetric multimodal fusion technique mCCA + jICA to uncover structural change patterns associated with FCPs using FNC matrices and GMV data from the ABCD dataset, comprising over 11,000 adolescent subjects across multiple scans. For each subject, we compute cell-wise ΔFNC and ΔGMV matrices, followed by estimating covarying multivariate patterns. Without predefined restrictions to specific seed regions, we calculate voxel-wise correlations between functional data loading parameters and ΔGMV data. The second approach involves using the mCCA+jICA multimodal fusion method to estimate covarying multivariate patterns FCPs and structural change patterns (SCPs).

Through both symmetric and asymmetric multimodal fusion techniques, our analysis identifies FNC linked to GMV data, suggesting concurrent changes between functional connectivity and structural data during adolescence. Notably, sex-related differences reveal stronger coupling between brain functional and structural changes in females compared to males. Our statistical analysis unveils several FCPs, and SCPs associated with longitudinal changes in psychopathology and cognition scores within the developing brain. The remainder of the research paper is organized as follows: in the Materials and Methods section, we present the data preprocessing and analysis procedures. In the Results section, we showcase changes in brain functional and structural coupling with age and the connections between various FCPs and SCPs with sex, differences in psychopathology and cognition scores. Finally, in the Discussion and Conclusion section, we delve into the implications of our findings.

## 2. Materials and Methods

### 2.1. ABCD DATA SUMMARY

The study utilizes data collected by the ABCD study ^1^. The primary objective of the ABCD study is to monitor brain development during adolescence. To do this the study collected a diverse array of data to discern the influences of biological and environmental factors on developmental trajectories. The ABCD study is the largest long-term investigation of brain development and child health in the United States. It includes multi-session MRI scans from over 11,800 children aged 9 to 11 years at baseline. The dataset covers subject details such as social, emotional, and cognitive development, gender identity, physical and mental health assessments, and medical backgrounds. Ethical considerations were upheld through parental informed consent and child assent, all approved by the Institutional Review Board (IRB). The ABCD dataset is accessible via the National Institute of Mental Health Data Archive (NDA) ^2^, where it has been made available as an open-source resource following its compilation from a diverse range of research endeavors across various scientific domains. The data were collected from 21 sites across the United States, ensuring standardized imaging methods. The quality of data was maintained through standard fMRI preprocessing and the Neuromark framework, a fully automated independent component analysis (ICA)-based approach that identifies brain networks across subjects [23]. The present study utilized data from 2,734 subjects who have both baseline and two-year follow-up scanned data of both the FNC and gray matter data. In our analysis, we specifically focused on the 1st scan from both the baseline and two-year data.

### 2.2. IMAGE PREPROCESSING OF FMRI DATA

We conducted preprocessing on the FastTrack fMRI images using a combination of the FMRIB Software Library v6.0 (FSL) toolbox and Statistical Parametric Mapping 12 (SPM) toolbox within the MATLAB 2019b environment. Initially, we corrected for rigid body motion by employing the mcflirt tool in FSL. Then, we conducted distortion correction utilizing fMRI field map data that were collected with phase-reversed blips. This process generated pairs of images with distortion occurring in opposing directions. To estimate the susceptibility-induced off-resonance field, we employed the FSL tool topup, utilizing volumes acquired with phase encoding both in the anteriorposterior and posterior-anterior directions. The coefficients derived from the output field map were then applied to correct distortion in the fMRI volume using the FSL tool applytopup. In the following step, we removed the initial ten scans with substantial signal changes to allow the tissue to stabilize in terms of radio frequency excitation. Subsequently, we spatially aligned the fMRI data to the standard Montreal Neurological Institute (MNI) space, utilizing the EPI template and resampling the data to 3 × 3 × 3 mm isotropic voxels via the spatial normalization tool in SPM. Lastly, we applied Gaussian smoothing with a full width at half maximum (FWHM) of 6 mm to the resliced fMRI images.

### 2.3. QUALITY CONTROL (QC)

We conducted data quality control (QC) on the preprocessed fMRI images to select subject data for subsequent analysis. The quality of how well subjects’ data were normalized to the MNI space has a significant impact on both the results of ICA and the estimation of FNC. Consequently, we excluded scans that did not exhibit satisfactory normalization to the MNI standard space. To be more specific, we compared individual masks with a group mask and retained scans that demonstrated strong similarities between their individual masks and the group mask. To achieve this, we initially calculated an individual mask for each scan of each subject based on the first fMRI time volume. Voxels were set to 1 if they exceeded 90% of the mean signal across the entire brain. Subsequently, we generated a group mask by designating voxels as 1 if they had more than 90% agreement with individual masks across the scans. For each scan, we computed spatial correlations between the group mask and the individual mask, focusing on the top 10 slices, bottom 10 slices, and the entire mask. This resulted in three correlation values for each scan. We included scans for further analysis if they met the following criteria: top-10-slices correlation greater than 0.75, bottom-10-slices correlation exceeding 0.55, and whole-brain correlation surpassing 0.8. This method ensures the inclusion of high-quality masks and fMRI data for retained scans, building on its success in previous research.

### 2.4. NEUROMARK FRAMEWORK

In this study, we used the Neuromark fMRI 1.0 network templates to extract intrinsic connectivity networks (ICNs) and their corresponding time courses (TCs) across subjects, via a fully automated spatially constrained ICA approach. The Neuromark fMRI 1.0 templates/priors were derived based on replicated networks estimated from two healthy control datasets, the human connectome project (HCP, 823 subjects after subject selection) and the genomics superstructure project (GSP, 1005 subjects after subject selection). Details of the Neuromark framework and templates can be found in the GIFT toolbox ^3^ and at ^4^ [23]. The selected spatial priors have also been demonstrated to be highly reliable between pipelines and across adult and adolescent datasets and populations [24]. Applying this approach yields 53 intrinsic connectivity networks for each subject. The resulting ICNs are highly corresponding and comparable across subjects, sessions, and scans. Children”s data can contain additional confounding effects such as larger head motions. To address this, we included four additional post-processing steps to control the remaining noise in the TCs of ICNs. The steps included: 1) detrend linear, quadratic, and cubic trends, 2) remove detected outliers, 3) apply multiple regression to remove variance linked to head motion parameters (3 rotations and 3 translations) and their temporal derivatives, 4) bandpass filtering using a cutoff frequency of 0.01 Hz-0.15 Hz. After the post-processing, we calculated Pearson correlation coefficients between post-processed TCs to estimate the static FNC for each scan.

### 2.5. PREPROCESSING OF SMRI DATA

We conducted preprocessing on the sMRI data using statistical parametric mapping^5^ within the MATLAB 2020b environment. The structural images were subjected to segmentation into gray matter, white matter, and CSF with additional modulation by the Jacobian to produce voxel-wise gray matter volume (GMV) maps. Subsequently, the GMV maps underwent smoothing via a Gaussian kernel with a FWHM of 6 mm.

### 2.6. ANALYSIS OF LONGITUDINAL CHANGE PATTERNS IN FNC AND GMV

In our research, we utilized subject-specific fMRI and sMRI data acquired during both the baseline and two-year scans to investigate changes in FNC and GMV. We calculated the cell-wise differences between the baseline and two-year FNC and GMV data to create ΔFNC and ΔGMV matrices respectively, representing the changes in FNC and GMV over time. These matrices were then analyzed using both asymmetric and symmetric fusion approaches to find link between brain functional connectivity and structure. In asymmetric fusion approach, we applied the ICA using the infomax algorithm [25] to deconstruct the ΔFNC in order to recognize longitudinal brain functional coupling and capture covarying patterns of changes, which are called FCP. We extended our analysis to investigate the connection between FCPs and brain structure. This involved calculating the voxel-wise correlation between the raw ΔGMV data and the loading parameters obtained from functional data after ICA estimation. More precisely, the second-level ICA model equation can be expressed as:

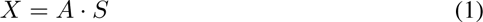

The following effectively represents the functional input data for the ICA model as:

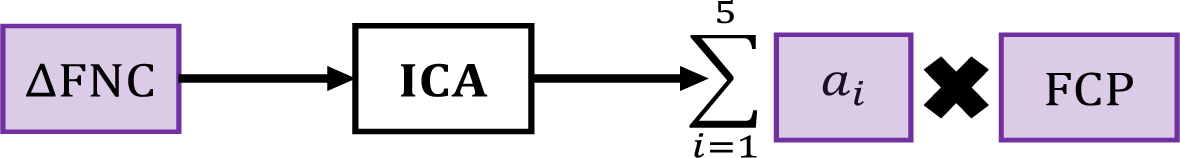

Next, we applied a symmetric fusion approach via mCCA+jICA to estimate joint SCPs and FCPs. In symmetric fusion approach, we run mCCA+jICA method to deconstruct the ΔFNC and ΔGMV matrices and identify patterns of change, namely functional and structural change patterns for FNC and GMV, respectively. We determined the optimal number of components, selecting five components for both GMV and FNC data using the elbow criteria. The mCCA+jICA model equation used in our experiment is expressed as follows:

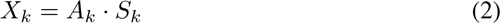

In this analysis, the dimensionality of the data matrix X is 2734 (subjects) × cells (for fMRI, the upper triangular elements of the ΔFNC matrix; for sMRI, the number of voxels). The dimensions of A are 2734 × 5 (components), S is 5 (components) × cells, and we have k = 2 is the number of modalities for the mCCA+jICA model.

This effectively represents the input data for the mCCA+jICA approach as following:

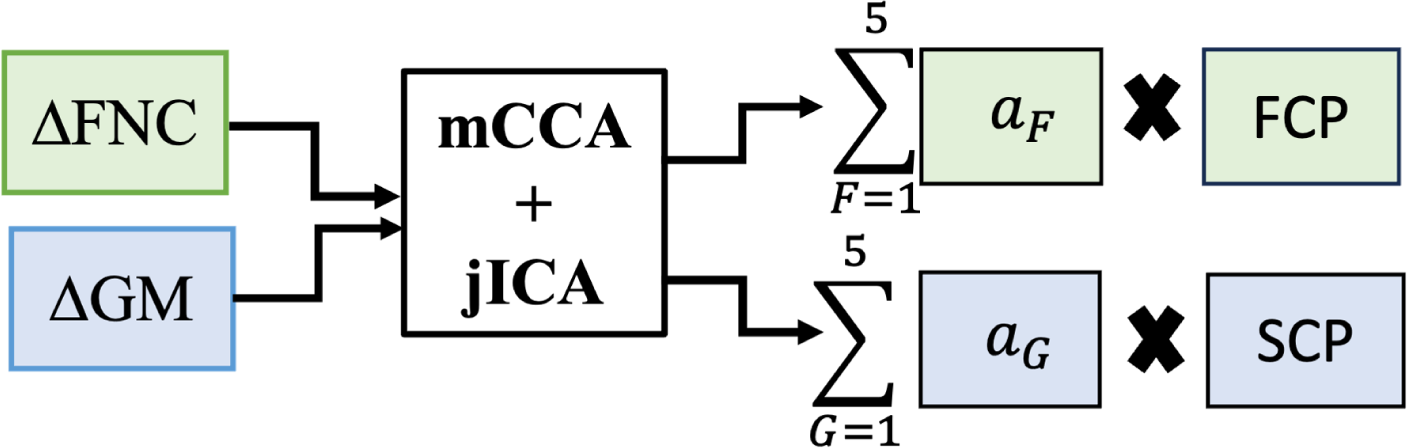

Here, ΔFNC represents the difference between the baseline (*F*_0_) and two-year (*F*_2_) functional network connectivity data, while ΔGMV corresponds to the difference between the baseline (*G*_0_) and two-year (*G*_2_) gray matter data. The FCPs and SCPs source matrices capture the most independent patterns of functional and structural changes, respectively. The parameter *a_i_* is each component’s subject-specific loading parameters of FNC data in the ICA model. The terms *a_F_* and *a_G_* denote the subject-specific loading parameters for each component in the FNC and GMV data, respectively in the context of the mCCA+jICA approach. These loading parameters quantify the individual subject’s contribution to the respective components. A block diagram of the analysis workflow is shown in figure 1.

**Figure 1:**
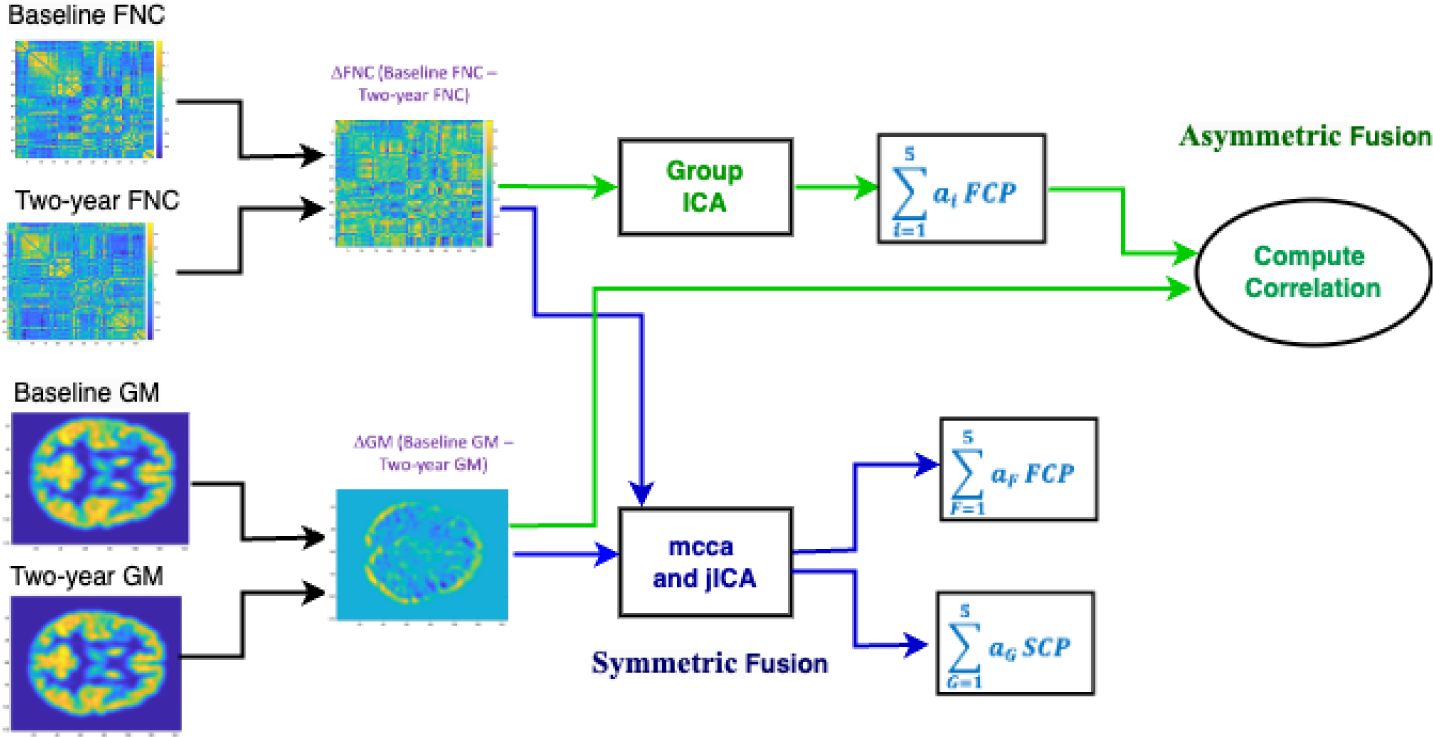
Block diagram of longitudinal multimodal functional and structural change pattern coupling analysis workflow. The subject-wise changed FNC and GMV matrix has been constructed by taking the difference between preprocessed baseline and two-year FNC and GMV data.

Moreover, following the mCCA+jICA estimation, we proceeded to evaluate the loading parameters and source matrix. In order to identify FCPs and SCPs that showed significant longitudinal changes compared to zero, we conducted one-sample t-tests on the loading parameters *a_F_* and *a_G_* for both modalities. We also conducted a one-sample t-test on the loading parameters (*a_i_*) following the ICA estimation. Statistical significance was assessed at a 95% confidence level, with adjustments made for multiple comparisons using a false discovery rate approach.

### 2.7. SEX BASED MULTIMODAL FUSION ANALYSIS

For the asymmetric multimodal fusion approach, we calculated correlations between ΔGMV data and loading parameters of functional data separately for males and females to examine sex differences in coupling. The next step involved computing the difference between these male and female correlations to understand the sex effect. In the symmetric multimodal fusion approach, we segregated the male and female loadings for both GMV and FNC data. Subsequently, we computed correlations between GMV male loadings and FNC male loadings, and between GMV female loadings and FNC female loadings. By quantifying the difference between these correlations (female - male), we assessed the strength of coupling between GMV and FNC loadings relative to sex. In both approaches, a significant positive difference would indicate stronger coupling between functional and structural data in females compared to males.

### 2.8. QUARTILE ANALYSIS OF PATTERN CHANGES VIA RELATIONSHIP TO SUBJECT MEASURES

In order to reduce the number of comparisons, we calculated composite scores for both cognitive and psychopathology assessments [26, 27]. Specifically, we computed the subject-wise differences between baseline and two-year data for composite psychopathology and cognitive scores, representing changes in psychopathology and cognitive performance, respectively. To explore the relationship among subjects who displayed the most significant age-related change pattern association with structural data, we conducted an analysis of the connection between loading parameters of ICA and ΔGMV data. We specifically selected subjects whose changes in psychopathology scores fell within or below the lower quartile. We then computed the correlation between the loading parameters of FCPs and the raw ΔGMV data for these chosen subjects, creating a voxel-wise association map for the lower quartile. Following the same procedure for subjects in the upper quartile, we also generated an upper quartile voxel-wise association map. Finally, we calculated the disparity between the upper and lower quartile voxel-wise correlation maps. The same methodology was applied to evaluate differences in cognitive scores. It’s important to note that all our findings underwent correction for multiple comparisons using the false discovery rate (FDR) [28].

## 3. Results

### 3.1. STRUCTURE-FUNCTION COUPLING STRENGTHENS WITH AGE ACROSS DEVELOPNEMT

The Neuromark fMRI 1.0 template encompasses a total of 53 reproducible networks, categorized into 7 domains based on their anatomical and functional attributes. These domains include subcortical, auditory, sensorimotor, visual, cognitive control, default mode, and cerebellar domains [23]. The experimental outcomes from the asymmetric fusion approach, involving spatial maps illustrating the links between multivariate FCPs and voxel-wise GMV data, are presented in Figure 2. In this figure, five FCPs components are plotted along with the spatial maps showcasing the voxel-wise correlations between GMV data and FCP loadings. Our symmetric fusion approach results, depicted in Figure 3, exhibit spatial maps illustrating the connections between multivariate FCPs and SCPs. This figure showcases five FCP components and their corresponding spatial maps of SCP components. Furthermore, the associations of FCPs and SCPs components with age are depicted using upper and lower arrows along with their associated T-values. A high negative (or positive) T-value signifies an increase (or decrease) in the expression of the specific change pattern with age [29], where the upper and lower arrow represent increasing and decreasing pattern changes with age, respectively.

**Figure 2:**
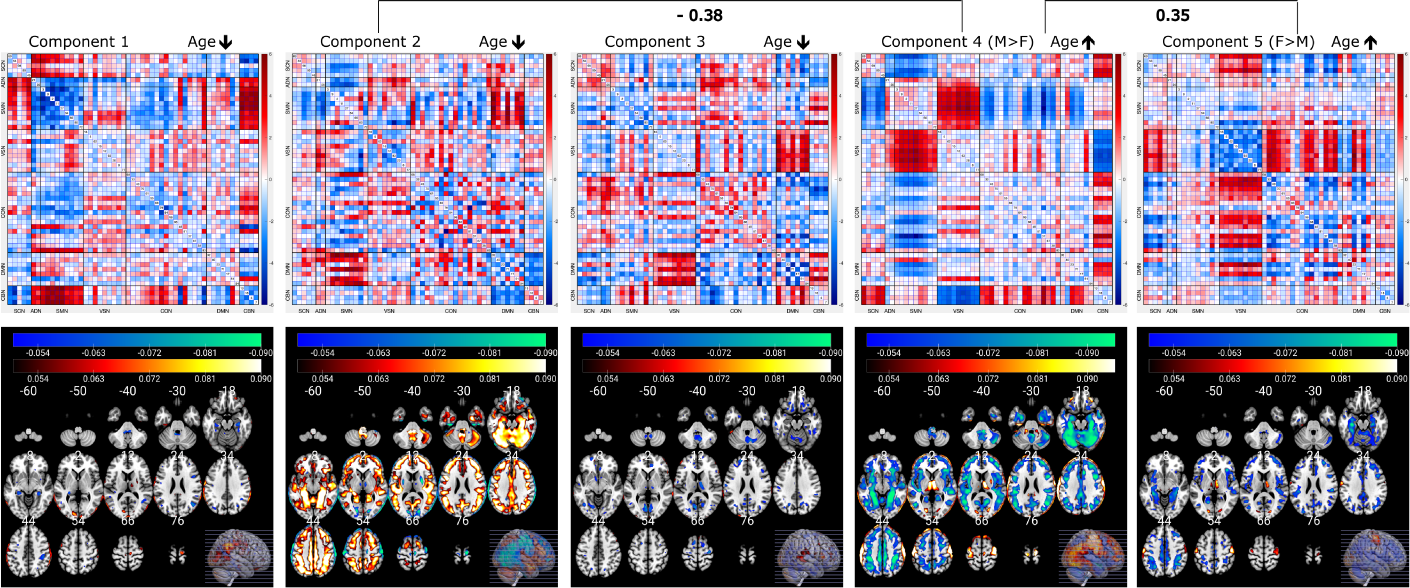
FNC component plots are displayed on the top, along with spatial maps depicting voxel-wise GMV correlation with the loading parameters of the FCPs are shown at the bottom for each component (asymmetric fusion). In the figure, we observe the voxel-wise correlation for components 2 and 4 have the highest positive (component 2) and negative (component 4) values.

**Figure 3:**
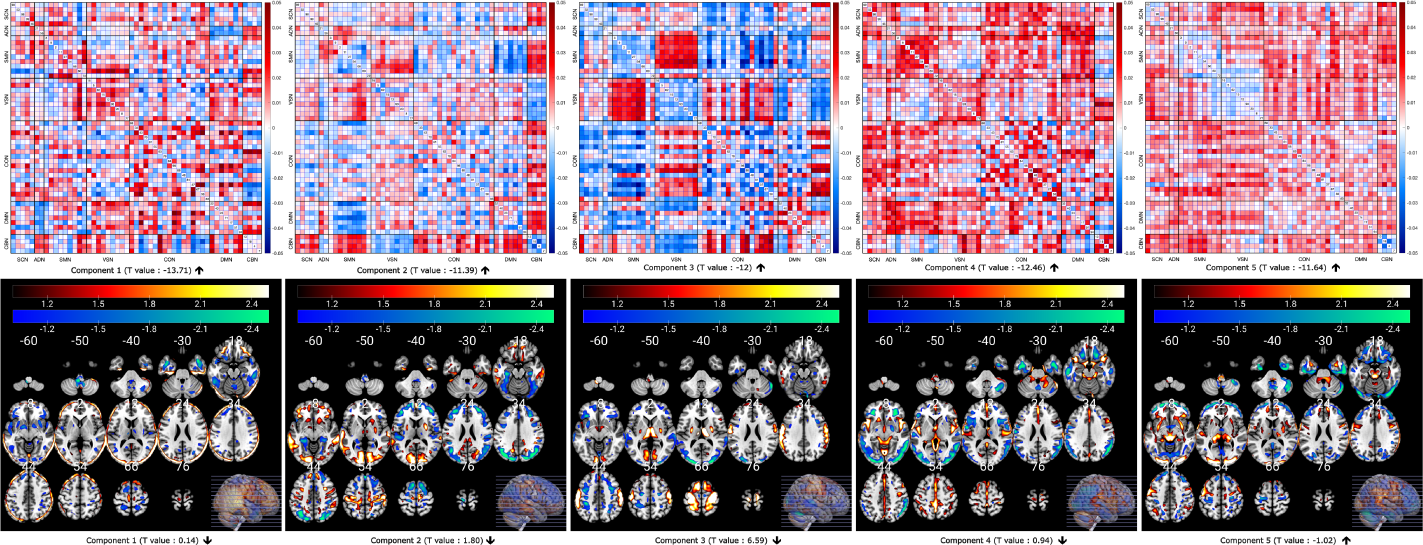
FNC components and spatial map of GMV components for the mCCA+jICA symmetric fusion approach. Here, component 3 from both functional and structural data exhibits highly structured changed patterns. In the figure, we observe increased functional connectivity between VS and SM domains or decreased functional connectivity between SM and CC correlated with SCPs indicating alterations in the bilateral sensorimotor cortex.

The results from the asymmetric fusion approach reveal noteworthy modularity in the fMRI result, suggesting structured changes occurring over the two-year period. And, interestingly, these were linked to strong and unique patterns of structural changes. In Figure 2, significant connections between FCPs and voxel-wise GMV data are evident. We also explored the covariation between the loading parameters. Notably, the correlation between loadings 4 and 2 of FCPs is -0.38. In the spatial map figure, component 2 displays a significant positive correlation (FDR-corrected, p = 2.45x-06), while the subcortical domain (SC) shows a widespread negative correlation with altered GMV data. The voxel-wise correlation map suggests that components 2 and 4 of FCPs exhibit the same correlation with altered GMV data, but in opposite directions. Furthermore, a strong positive correlation (r = 0.35) between loadings 4 and 5 of FCPs is observed. Similar to component 4, our voxel-wise correlation map reveals a significant negative correlation between FCPs and voxel-wise changed GMV data for component 5 (FDR-corrected, p = 9.83x-04). Conversely, the SC shows a significant positive correlation, with notable differences in the subcortical region of the voxelwise correlation between components 2 and 4. Component 4 demonstrates a higher voxel-wise correlation in the SC.

The results from the symmetric fusion approach indicate that FCPs associated with components 2 and 3 exhibit significant changes with increasing age in the developing brain. Both components demonstrate an increasing trend with age, as evidenced by their negative T-values. Component 3 reveals an increased brain functional connectivity between the visual domain (VS) and sensorimotor domain (SM) in the FNC data. Correspondingly, there are decreasing changes in the bilateral sensorimotor cortex in the sMRI data over the two-year period. Additionally, the FCP of component 3 demonstrates a decreasing trend with age in the functional connectivity between the VS and cerebellar domain (CB), as well as between the SM and cognitive control domain (CC). Additionally, we conducted a two-sample t-test based on sex information using the loading parameters of both modalities. Our analysis revealed that males exhibit smaller change pattern expression in SCP for component 2 compared to females.

### 3.2. EVALUATION OF GENDER SHOWING CHANGE PATTERNS COUPLING

In asymmetric multimodal fusion method, we have computed the difference (female-male) in the voxel-wise correlation map between females and males to assess the sex effect. We found a positive correlation between the loading parameter of component 2 and raw ΔGMV for both male and female data. Component 4 shows negative association with ΔGMV and significant correlation difference between males and females where females show higher voxel-wise correlation than males as displayed in figure 4. We conducted FDR correction on the correlation values with a significance threshold set at 0.001.

**Figure 4:**
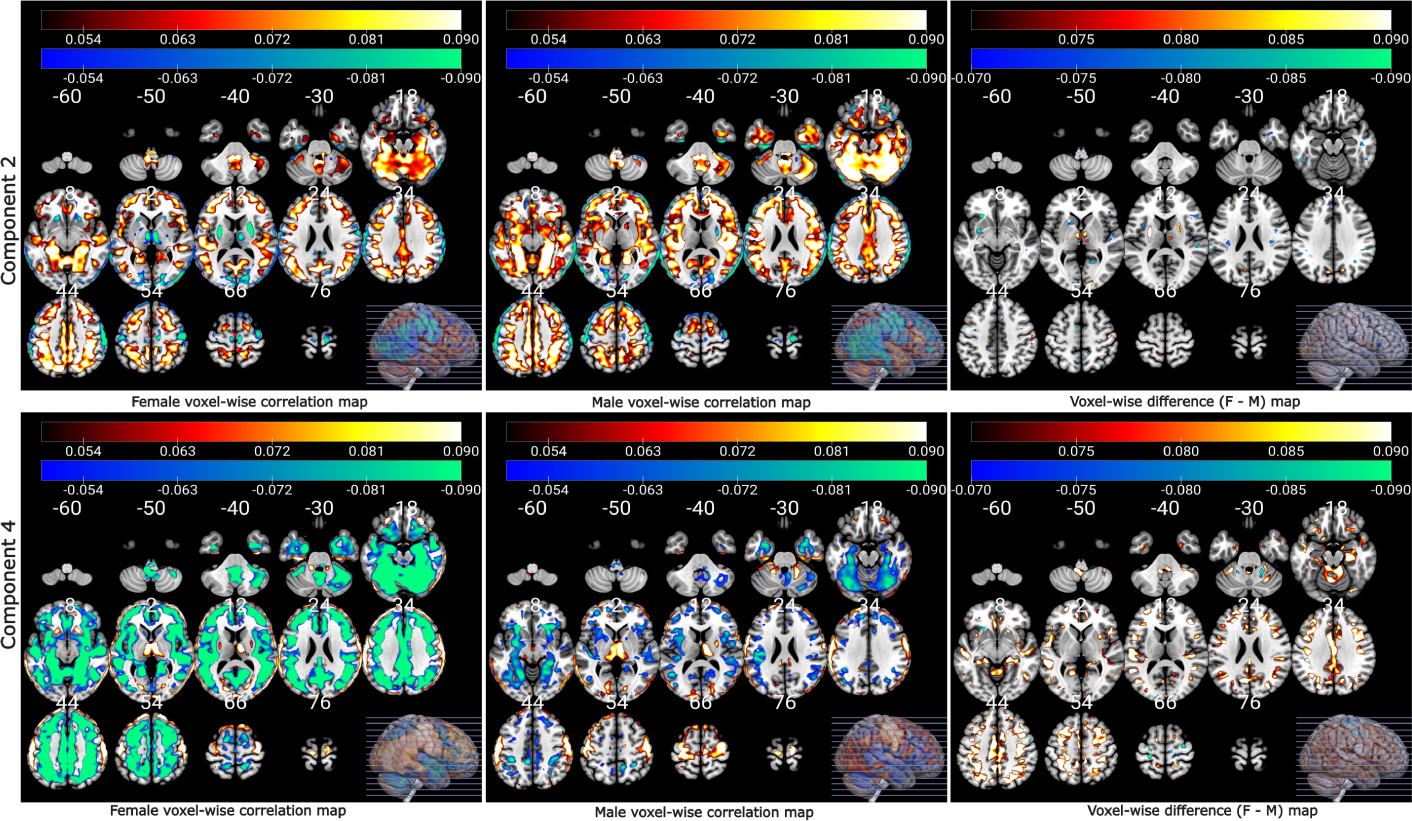
Voxel-wise correlations of females, males, and distinctions in voxel-wise correlations map (female - male) between male and female loading parameters for FCPs and raw ΔGMV. Females exhibit a stronger positive coupling for the FCP component 2 and stronger negative coupling for the FCP component 4 compared to males.

After applying the symmetric multimodal fusion technique, we computed the Pearson correlation between the loading parameters for all FCPs and SCPs, separately for males and females. This analysis aimed to explore sex differences in coupling, specifically the associations at the subject expression level. The variations in correlations between loading parameters of FCPs and SCPs for males and females are visualized in Figure 4. Our findings indicate that females exhibited stronger coupling between SCP component 2 and FCP component 1 (Δ r = 0.128, FDR-corrected, P = 2.1895e-11), FCP component 3 (Δr = 0.102, FDR-corrected, P = 1.0081e-07), and FCP component 4 (Δr = 0.111, FDR-corrected, P = 6.7136e-09) compared to males.

### 3.3. ASSOCIATION OF CHANGES IN MULTIVARIATE PATTERN COUPLING WITH PSYCHOPATHOLOGY SCORE, AND COGNITIVE SCORE

We explored the relationship between changes in multivariate pattern coupling and subject measures, specifically psychopathology scores difference and cognitive scores difference. Our results reveal that both components 4 and 5 exhibit a significant negative association with raw ΔGMV data within the lower quartile of psychiatric scores, as illustrated in Figure 6. The voxel-wise correlation maps for cognitive scores are presented in Figure 7. Component 2 of FCP predominantly shows a positive correlation with the ΔGMV data within the lower quartile of cognitive scores. Regarding component 5, it displays negative correlations in the lower quartiles. We observed a noteworthy difference between the upper and lower quartiles for both psychopathology and cognitive scores. Furthermore, we applied FDR correction at a significance level of 0.05 to the correlation values and only significant results are shown in the figure.

**Figure 5:**
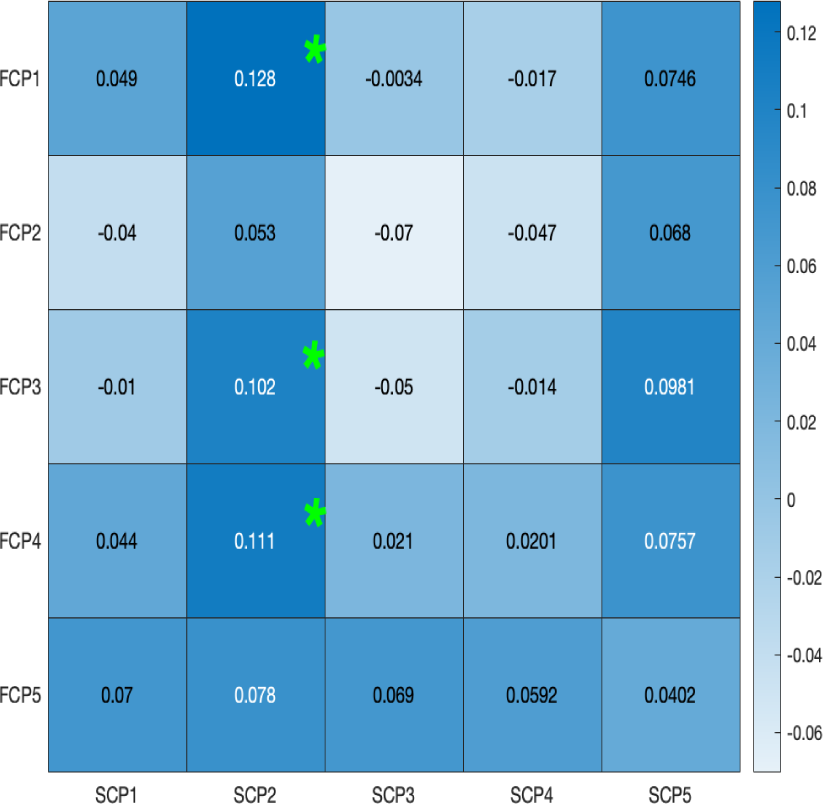
Differences in correlations between male and female loading parameters for FCPs and SCPs. Females exhibit a stronger coupling between the FCP component 1, 3 and 4 and SCP component 2 (indicated by *) expressions compared to males.

**Figure 6:**
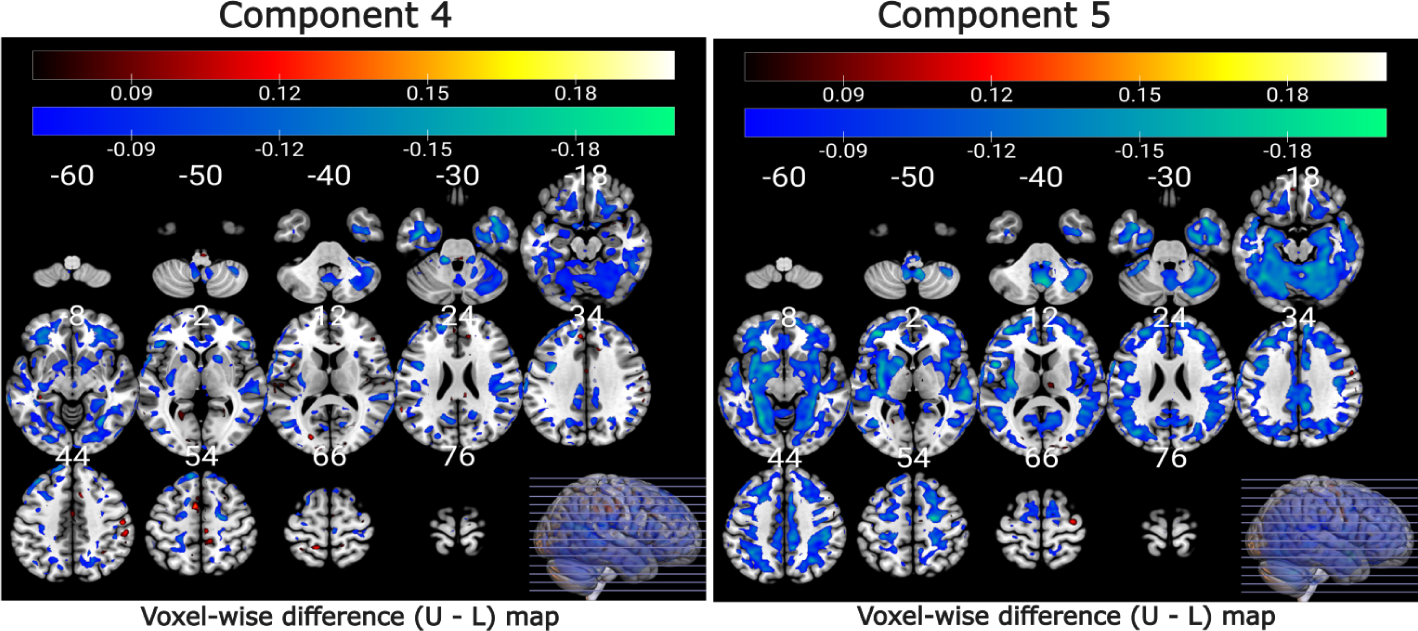
Voxel-wise correlation difference map (upper - lower) illustrating the relationship between loading parameters of FCPs and raw ΔGMV for the upper and lower quartile of psychiatric scores. Components 4 and 5 exhibit notably stronger voxel-wise correlations within the lower quartile of psychiatric scores (differences are negative).

**Figure 7:**
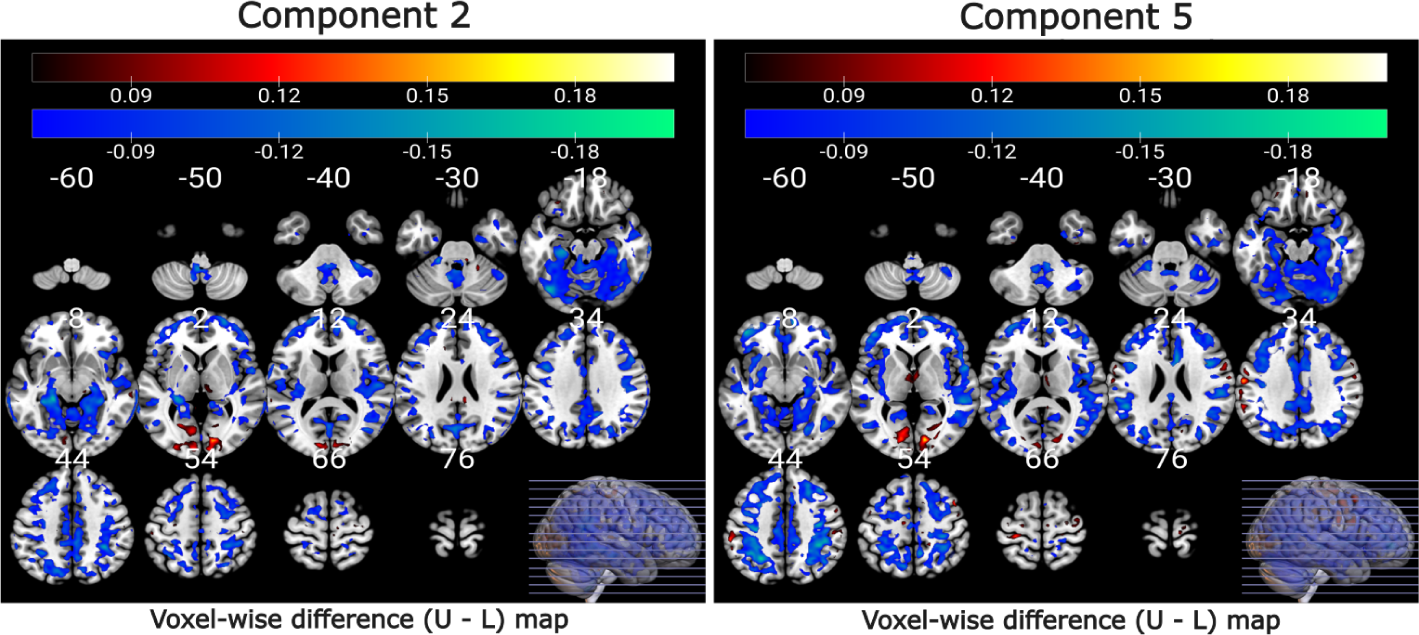
Voxel-wise correlation difference map (upper - lower) illustrating the relationship between loading parameters of FCPs and raw ΔGMV for the upper and lower quartile of cognitive scores. Components 2 and 5 exhibit notably stronger voxel-wise correlations within the lower quartile of cognitive scores

## 4. Discussion

To the best of our knowledge, this is the first study to estimate the multivariate pattern coupling/covariations in structural and functional connectivity associated with age progression using a very large sample of nearly 2734 subjects in developing brain. Our study introduces two data-driven multivariate multimodal fusion analyses to delineate change patterns in brain structure and function among healthy adolescents over a two-year period. The key findings include 1) identification of intriguing patterns in FNC and GMV data, demonstrating links to individual variations. Specifically, FCP components 2 and 4 display significant associations between their loadings and voxel-wise GMV data, 2) discovery of structured FCPs, such as enhanced connectivity between visual and sensorimotor domains or decreasing coupling trend between the SM and CC, correlated with SCPs indicating alterations in the bilateral sensorimotor cortex, 3) observation of stronger coupling between brain functional and structural changes in females compared to males, unveiling sex-related disparities, 4) recognition of multiple FCPs and SCPs linked to longitudinal changes in psychopathology and cognition scores within the developing brain.

Understanding longitudinal changes in the developing brain is crucial for comprehending the dynamic processes underlying cognitive development and neural maturation. The integration of functional connectivity and sMRI through multimodal fusion offers a powerful avenue to explore these intricate developmental patterns. The results of this research underscore the intricate relationship between functional and structural changes occurring over time. By employing multimodal fusion techniques, significant connections between FCPs and voxel-wise GMV data have been revealed. This investigation aligns with previous studies emphasizing the significance of multimodal imaging techniques in comprehending the complex dynamics of brain maturation [30, 31]. By leveraging both FCPs and voxel-wise GMV data, the study brings to light notable findings. The utilization of asymmetric fusion unveils compelling modularity trends over a two-year period, suggesting structured changes occurring during brain development [32]. The identification of significant connections between FCPs and voxel-wise GMV data corroborates the interplay between functional and structural aspects of brain maturation, echoing earlier studies emphasizing the association between functional connectivity and anatomical substrates [33, 34, 35]. The observed correlations between specific FCP loadings unveil complex relationships between functional connectivity patterns and altered GMV data. These findings suggest that changes in functional connectivity are associated with corresponding alterations in brain structure, indicating a tightly intertwined relationship between brain function and structure during development. The correlation analysis between loading parameters sheds light on intriguing relationships within FCPs. Notably, the negative correlation between specific loadings, such as the -0.38 correlation between loadings 4 and 2, emphasizes the complexity of brain development and the potential opposing effects within functional connectivity networks [36]. The observed patterns, such as components 2 and 4 exhibiting similar but opposite correlations with altered GMV data, underscore the nuanced and region-specific nature of brain development [37]. These findings resonate with the growing body of research aiming to decipher the intricate interplay between functional and structural changes in the developing brain. They invite further exploration into the specific mechanisms governing the observed modularity and correlations, potentially offering deeper insights into the underlying processes of brain maturation. Additionally, the spatial mapping of components reveals differential correlations with altered GMV data, particularly in the SC domain. The observed patterns in the voxel-wise correlation map highlight regional specificity in the relationship between FCPs and altered GMV data. Notably, the contrasting correlations in different brain regions emphasize the importance of considering regional variations in brain development. Such insights contribute to our understanding of how specific brain regions may exhibit distinct developmental trajectories, potentially influencing cognitive and behavioral outcomes.

The utilization of the symmetric multimodal fusion approach, mCCA+jICA reveals intricate relationships between structural and functional changes over time. Numerous prior studies [16, 38, 39, 5, 33, 40, 41, 42, 43, 44, 45] distinctly indicate associations between structural and functional connectivity. The findings showcase co-varying patterns in longitudinal changes, shedding light on the dynamic interplay between structural alterations and functional connectivity in the maturing brain. These observations echo previous research highlighting the mutual influence and interaction between brain structure and function during developmental stages [46, 30]. Our analysis reveals that both component 2 and 3’s FCPs have negative T-values, indicating increasing trend with age in adolescents. For component 3, we observed an increasing functional connectivity coupling between the VS and SM domain over the two-year period. The interaction between the VS and the sensorimotor SM network is crucial for integrating visual and motor information to support perception, action, and cognition. This interaction enables the accurate and coordinated execution of movements, the integration of sensory and visual feedback, and the control of attention and working memory [47]. This evidence indicates that the developing brain begins to establish connections among different regions, while simultaneously adapting its neural and cognitive systems to respond to external stimuli. It is reasonable to infer that these connections among neural circuitry in the developing brain play a critical role in creating distinctions between regular and irregular neural dynamics. The observed enhancement in connectivity between the visual and sensorimotor domains is consistent with previous research emphasizing the dynamic interconnectivity between these areas. Studies by Dayan and Cohen [48] and Ungerleider and Haxby [49] have highlighted the relevance of visual inputs in motor coordination and sensorimotor integration. We also observed a decreasing functional coupling between the VS and CB and SM and CC regions with age, which exhibited the largest positive T-values. The findings of reduced coupling between domains in this study potentially indicate a shift in the coordination or interaction between these brain regions, reflecting a reorganization of neural networks. The correlation of these identified FCP alterations with SCPs indicating alterations in the bilateral sensorimotor cortex strengthens the relationship between functional connectivity and underlying structural modifications. The results highlight instances where structural changes are accompanied by corresponding alterations in functional connectivity patterns. Such co-varying patterns observed in the brain’s structural and functional dynamics imply a close relationship between these modalities during development [39]. Additional studies suggest that alterations in neuronal functioning are directly related to alterations in gray matter structure [50]. The significance of these findings lies in their potential to uncover underlying mechanisms driving brain development and maturation. Understanding how structural changes co-vary with alterations in functional connectivity provides valuable insights into the coordinated and interdependent nature of brain development [51, 52]

In this study, we employed both asymmetric and symmetric multimodal fusion methods to explore sex effects on functional and structural change coupling, aiming to discern nuanced differences in neural dynamics between males and females. In the asymmetric multimodal fusion method, the examination of voxel-wise correlation maps revealed intriguing sex effects. Specifically, a positive correlation between the loading parameter of component 2 and raw ΔGMV was observed for both males and females. This finding is in line with previous studies indicating sex-related differences in brain structure and connectivity, suggesting that both sex exhibit similar patterns of association between specific components and structural changes [53]. However, component 4 exhibited a sex-specific association with ΔGMV, demonstrating a significant correlation difference between males and females. Females exhibited higher voxel-wise correlation than males, suggesting a nuanced sex-related divergence in the relationship between this specific FCP and structural changes. Sex-specific differences in brain structure and connectivity have been previously reported [54], and the present study adds granularity by highlighting component-specific distinctions. The application of the symmetric multimodal fusion technique further delved into sex differences in coupling at the subject expression level. The results revealed distinct variations in coupling patterns between males and females, shedding light on sex-dependent associations within the neural network. Notably, females exhibited stronger coupling between SCP component 2 and FCP components 1, 3, and 4 compared to males. This discovery indicates that women who play a major role in the structural change pattern (SCP component 2) also make substantial contributions to the FCPs of components 1, 3, and 4. These observations suggest sex-dependent variations in the interplay between structural changes and functional connectivity, emphasizing the importance of considering individual differences in understanding brain dynamics [53, 55]. Based on the obtained correlation values, we can conclude that females exhibit a stronger coupling between the function and structure of developing brain compared to males.

Our analysis uncovered several functional and structural connectivity change pattern coupling associated with longitudinal changes in psychopathology and cognition scores within the developing brain. Specifically, we observed a significantly higher negative correlation between FCP components 4 and 5 and raw ΔGMV in subjects with low psychiatric scores compared to those with high psychiatric scores. This finding implies that individuals with lower psychiatric scores demonstrate a stronger association between FCPs, such as increased brain functional coupling between the VS-SM and CB-SC, or a decrease in change patterns between VS-CB and SM-SC, and raw ΔGMV. Additionally, FCP components 2 and 5 exhibit positive and negative associations, respectively, with the ΔGMV data within the lower quartile of cognitive scores. This indicates that individuals with lower cognitive scores show both positive (component 2) and negative (component 5) voxel-wise correlations. Notably, a significant higher correlation is observed between both FCP components 2 and 5 and raw ΔGMV within the lower quartile of cognitive scores compared to the upper quartile. This suggests that individuals contributing less to changes in cognitive scores from baseline to two years exhibit a stronger association between functional and structural data in the developing brain. These findings align with recent studies that have reported age-related changes in vasculature, brain anatomy, and brain function collectively contributing to complex interactions that influence cognitive alterations [56, 57].

In this manuscript, we present an innovative method that combines various data types to analyze patterns across multiple domains. We apply this method to explore the connection between brain structure and function using a combination of FNC-sMRI data. Several promising research directions are apparent for further exploration. For instance, expanding the use of additional imaging techniques could enhance our understanding of overall brain change patterns, a path we aim to explore in future investigations. Furthermore, future research should delve into how increased component numbers affect these couplings, where varying the component count may reveal stronger connections between different brain regions. Our proposed methods also have certain constraints that warrant consideration. First, it’s important to note that while mCCA+jICA operates on ICA components instead of the original imaging data (e.g., using 3D contrast images instead of 4D fMRI data), this approach may lose some temporal information. However, working with these ICA components presents advantages such as reduced dimensionality [6] and a simplified space for linking the data [58]. Second, our utilization of ICA assumes linearity in capturing brain functional pattern changes. Nonetheless, recent research by Motlaghian et al. highlights the potential presence of nonlinear relationships within functional networks, a factor often overlooked in linear analyses [59]. Exploring nonlinear methodologies could offer valuable insights into age-related functional change patterns in the developing brain. Finally, this study utilized data from youths aged 9-10 years, potentially indicating relatively low levels of psychiatric symptomatology among the subjects [60]. However, it’s anticipated that their psychopathology load may increase during adolescence [61]. This relationship between brain and behavior might evolve growing stronger, weaker, or altering in some other way which could be directly examined in future waves of longitudinal ABCD data. This would enable the formulation of clear hypotheses.

## 5. Conclusion

In this study, we propose both asymmetric and symmetric multimodal fusion techniques to investigate longitudinal brain change patterns in adolescents. These methods are designed to explore comprehensive structural and functional changes and their correlations with aging, utilizing FNC matrices and GMV data from the ABCD dataset. Our results from the asymmetric multimodal fusion technique reveal significant associations between FCP components 2 and 4 and voxel-wise GMV data. We hypothesize that the multivariate patterns of FNC and GMV demonstrate relationships with increased age. Using the symmetric multimodal fusion technique, we identified structured FCPs, such as increased connectivity between visual and sensorimotor domains and decreasing connectivity between the sensorimotor and cognitive control, correlated with SCPs involving alterations in the bilateral sensorimotor cortex. Studying the coupling between functional connectivity and GMV over time aids in understanding how the developing brain modulates its structural and functional coupling and relationships with other brain regions for efficient information processing. Our experiments indicated specific brain components that significantly contributed to sex-based differences in brain functional and structural coupling across both multimodal fusion techniques. Additionally, we uncovered several functional and structural couplings associated with psychopathology and cognition in the developing brain. Importantly, our proposed methodologies for investigating age-related change patterns and coupling in the developing brain is the first approach that can be a useful tool to evaluate whole-brain co-varying functional and structural change patterns in longitudinal data

## ACKNOWLEDGEMENT

The work was funded by the NIH (R01MH123610) and the NSF (2112455).

## Footnotes

1 https://abcdstudy.org/

2 https://nda.nih.gov/

3 [http://trendscenter.org/software/gift

4 http://trendscenter.org/data

5 SPM12http://www.fil.ion.ucl.ac.uk/spm/

